# Global Metabolomic Analysis of Lytic KSHV Infection: Induced Host Nucleotide Metabolism is Required for Infectious Virus Production

**DOI:** 10.64898/2026.02.02.703314

**Authors:** Fatima Hisam, Emma Winn, Spandan Mukherjee, Savannah Price, Yennifer A. Gaspar, Claire Wang, Hamid R Baniasadi, Tracie Delgado, Erica L. Sanchez

## Abstract

Kaposi’s Sarcoma Herpes Virus (KSHV) is the etiological agent of Kaposi’s Sarcoma (KS) which is known to cause metabolic stress in infected host cells. KSHV reprograms host metabolic pathways for efficient viral replication and infectious virion production. Here, we report a time-course global metabolomics study conducted in the doxycycline-inducible iSLK.BAC16 cells to compare latent and lytic KSHV infection. Our data show that amino acid, central carbon, and nucleotide metabolic pathways are highly dysregulated upon reactivation to lytic replication. During lytic KSHV infection, pathway enrichment analysis shows that the top two most significantly impacted and dysregulated pathways are purine and pyrimidine metabolism. Further experiments have shown that nucleotide metabolism is required during lytic KSHV infection to produce maximal infectious virus. Treatment with the FDA-approved drug, methotrexate (MTX), a folate antagonist that inhibits cellular DHFR and decreases nucleotide metabolism by reducing tetrahydrofolate cofactors, significantly reduced KSHV late lytic viral gene expression upon reactivation compared to control. Additionally, titering cell-free supernatants from MTX-treated lytic KSHV-infected cells showed a significant reduction in infectious virion production. Furthermore, by adding folinic acid (FA), a downstream metabolite of the MTX-DHFR inhibition step, in the presence of MTX, late lytic gene expression and infectious virion production were significantly rescued. Furthermore, we observed a significant decrease in viral titer of murine herpesvirus 68 (MHV-68), a model virus to study gammaherpesvirus, after MTX treatment. Overall, our study demonstrates that metabolic inhibition during lytic gammaherpesvirus infection decreases productive infection and hence, serves as a potential therapeutic antiviral target.

## Introduction

Kaposi’s sarcoma-associated herpesvirus (KSHV), an oncogenic gammaherpesvirus, is the causative agent of Kaposi’s Sarcoma (KS) (1, 2). KS is a soft-tissue cancer that presents as red-purple lesions on the skin and mucous membranes. KSHV is also involved in other human malignancies, including primary effusion lymphoma (PEL), multicentric Castleman’s disease (MCD), and KSHV inflammatory cytokine syndrome (KICS) (1, 3). KSHV causes cancer in immunosuppressed individuals and is the most common cancer in AIDS patients across the globe (3, 4).

Like all herpesviruses, KSHV is a double-stranded DNA virus (5) enclosed in a protein capsid and then in a lipid bilayer (6). KSHV undergoes two distinct viral phases: latent and lytic infection. The majority of cells in KS lesions are latently infected, with approximately 5% undergoing lytic replication. The latent phase is defined by limited viral gene expression, including expression of the Latency Associated Nuclear Antigen (LANA). This stage results in no new virion production, and the host cell survives and proliferates (7). The lytic viral phase expresses all the genes in a temporal fashion which includes immediate early and early viral gene expression, viral genome replication, late viral gene expression, virion production, and egress from cell. Representative viral genes investigated in this study are the latent gene LANA, the early viral gene DNA polymerase (ORF59), and the late viral gene viral glycoprotein (K8.1) (8).

Viruses are obligate parasites that depend on the host cell for their growth and survival. Host cell metabolism is one of the major cellular systems manipulated during KSHV infection. KSHV, like other herpesviruses, commandeers host cell metabolism for energy production and raw material synthesis for viral replication and production (9, 10). However, the virus-host cell interactions and many of the underlying mechanisms of these viral manipulations of cellular metabolic pathways remain unclear.

The field of metabolomics has emerged as a key discipline in the wake of genomics, transcriptomics, and proteomics(11). Studies have been done to quantify the levels of a large number of metabolic compounds (energy molecules and biochemical building blocks) during viral infection in host cells by employing an advanced measurement technique, liquid chromatography-mass spectrometry (LC-MS)(12). Previous studies have shown that viral infections like Human Cytomegalovirus (HCMV), Epstein-Barr virus (EBV), Hepatitis C (HCV), Human Immunodeficiency Virus (HIV), Dengue, and Herpes Simplex Virus (HSV-1) lead to dramatic increases in the levels of many metabolites and that these increases substantially exceed those associated with normal transitions of cells between resting and growing states. In several studies, differential metabolite abundances induced by viral infection coincide with an increase in host cell expression of enzymes that drive metabolite synthesis (13–18)

Mass spectrometry-based metabolomic approaches confirmed the divergent metabolic profile of two related herpesviruses, HCMV and HSV-1(16). These viruses need different metabolites based on the duration of viral replicative cycles. HSV-1 has an average replication cycle of 24 hours and requires rapid accumulation of nucleotides, which is marked by a significantly increased demand for DNA synthesis. On the other hand, HCMV can accumulate nucleotides over time since it has a replication cycle of ∼96 hours, altering the glycolytic metabolites used in the production of fatty acid synthesis and inducing the production of new lipid membranes (16).

Previously, both gas chromatography-mass spectrometry (GC-MS) and/or Liquid chromatography-mass spectrometry (LC-MS) were performed on latent KSHV-infected endothelial cells compared to mock-infected controls, which demonstrated that significant metabolic pathways were altered by latent infection, similar to those altered in cancer cells (19). Latent KSHV infection is seen to have increased amino acid metabolism, including polyamine metabolism like putrescine, spermidine, and spermine, after 48 and 96 hours (19). KSHV latent infection alters glycolysis and induces the Warburg Effect in the infected cells, resulting in decreased glycolytic flux to oxidative phosphorylation and increased lactate production, a key hallmark of cancer (20, 21). KSHV latent infection also requires lipogenesis and increases lipid droplet formation for lipid storage, which is necessary for virion assembly (19, 21). The requirement of cellular metabolism for KSHV latent infection was further correlated with proteome and phosphoproteome analysis.(22) An untargeted metabolomics of KSHV latent infection versus mock determined the metabolic changes induced after four hours of latent infection, including urea cycle, polyamine synthesis, dimethylarginine synthesis, and de novo pyrimidine synthesis(23). A recent KSHV study explored mechanistic biochemistry and metabolic pathway manipulation, which focused on metabolic rewiring that supports latent cell proliferation. The study demonstrated that KSHV hijacks host nucleotide synthesis and glycolysis via carbamoyl-phosphate synthetase 2, aspartate transcarbamoylase, and dihydroorotase (CAD) and RelA, hence promoting viral persistence and inducing proliferation (24). Overall, these previous studies highlighted metabolic pathways implicated in KSHV infection; however, further exploration of host cell metabolic alterations during KSHV lytic replication are necessary.

Recently, metabolomics and lipidomics analysis performed on murine herpesvirus 68 (MHV-68), a mouse gammaherpesvirus which shares significant homology to human gammaherpesvirus KSHV and EBV, induces alteration in key metabolic pathways like glycolysis, glutaminolysis, lipid metabolism, and nucleotide metabolism(25). Interestingly, nucleotide metabolism was the highest increased metabolic pathway in the lytic MHV-68 metabolomics. Furthermore, inhibition of glycolysis and lipogenesis through pharmacological drugs and inhibition of glutaminolysis through starvation decreased infectious MHV68 virus production. These results suggest that lytic gammaherpesvirus infection is highly dependent on the alteration of key metabolic pathways in favor of infectious virion production. However, this had yet to be evaluated in a lytic model of KSHV infection.

Here, we report that KSHV viral replication is dependent on host nucleotide metabolism. The experiments were conducted using the iSLK.BAC16 cell line, which stably maintains the recombinant KSHV-BAC16 encoding constitutive expression of GFP (26). These cells encode the viral Replication and Transcription Activator (RTA) transgene and are doxycycline (Dox) and sodium butyrate (NaB) inducible, which triggers the transition from latent to lytic infection(26). LC-MS analysis of lytic KSHV infection in iSLK.BAC16 cells compared to latent infection revealed alteration in host metabolic pathways, including amino acid metabolism, central carbon pathways, and nucleotide metabolism. Metabolic pathway enrichment analysis revealed the top two most significantly impacted and dysregulated pathways to be purine and pyrimidine metabolism. In our LC-MS data, the increased abundances in key purine/pyrimidine-triphosphate species during lytic infection suggest an amplified shift in nucleotide biosynthesis, potentially to support the production of more viral genes and transcripts to sustain infection. Inhibition of pyrimidine and purine synthesis via the FDA-approved drug methotrexate (MTX) significantly reduced extracellular viral titers of KSHV and MHV-68. MTX, a competitive inhibitor of dihydrofolate reductase (DHFR), is a folate antagonist that is used as a chemotherapeutic agent and immunosuppressive drug inhibitor (27–29). MTX functions by decreasing tetrahydrofolate (THF), which acts as a substrate for subsequent metabolic steps (30, 31). Interestingly, by rescuing the toxicity of MTX with folinic acid (FA, 5-formyl THF), we see an increase in extracellular virion production. Additionally, MTX treatment on iSLK.BAC16 cells demonstrated a significant reduction in late lytic viral mRNA expression while supplementing FA to lytic KSHV infected cells in the presence of MTX, we see an increased expression in late lytic viral mRNA expression.

Our global metabolomic analysis reveals the changes in key metabolites of central carbon pathways, amino acid metabolism, and nucleotide metabolism during the infection time course after reactivation. Furthermore, we demonstrate that both KSHV and MHV-68 lytic infections require functional nucleotide metabolism for maximal viral infection. Understanding how KSHV lytic infection drives virus-induced global host cell metabolic changes and recapitulating the importance of altered host nucleotide metabolism during MHV-68 lytic infection can provide us with insights into new therapeutic mechanisms to manage KSHV-associated malignancies.

## Results

### KSHV lytic infection drives global metabolic alterations

iSLK.BAC16 cells stably maintaining the KSHV genome were used to determine the lytic viral metabolic profiles compared to latent infection at three different time points post Dox/NaB mediated induction. Metabolomic analysis was performed at 24, 36, and 48 hours post-induction (hpi) via Dox and NaB (**Figure 1A**). The targeted LC-MS analysis included five biological replicates normalized to protein concentration measured via BCA. A total of 160 metabolites were detected across samples: 56 metabolites for amino acid metabolism, 61 metabolites for central carbon metabolism, and 44 metabolites for nucleotide metabolism (**Supplemental Table 1**). Principal Component Analysis (PCA) was conducted in MetaboAnalyst 6.0 to visualize global metabolic variation across samples. The score plot displaying each sample according to its principal components (PC), revealed natural clustering patterns and differences in metabolic profiles between latent and lytic samples at different time points (**Figure 1B**). Separation along PC1 or PC2 indicated the consistent metabolic shifts associated with the various experimental conditions, while overlap indicates similarity between the sample sets (**Figure 1B**). One-way ANOVA was performed on the latent and lytic samples across different time points with a false discovery rate (FDR) cutoff at 0.05, and the data was subjected to Tukey’s Honestly Significant Difference (HSD) post-hoc analysis. The scatter plot displayed the significant difference in the abundances of metabolites across samples. Among 160 detected metabolites, 143 metabolites exhibited statistically significant differences **(Figure 1C, Supplemental Table 2)**. The y-axis corresponds to -log10(raw p-value), and the x-axis corresponds to the metabolites. The top discriminating metabolites include the highest FDR values, indicating strong between-group differences. Pathway enrichment analysis reveals the top two most significantly impacted and dysregulated pathways to be purine and pyrimidine metabolism (**Figure 1D**). Other pathways in the top 25 dysregulated pathways are amino acid metabolism, the citrate cycle, glutathione, arginine, and thiamine metabolism.

**Figure 1.**
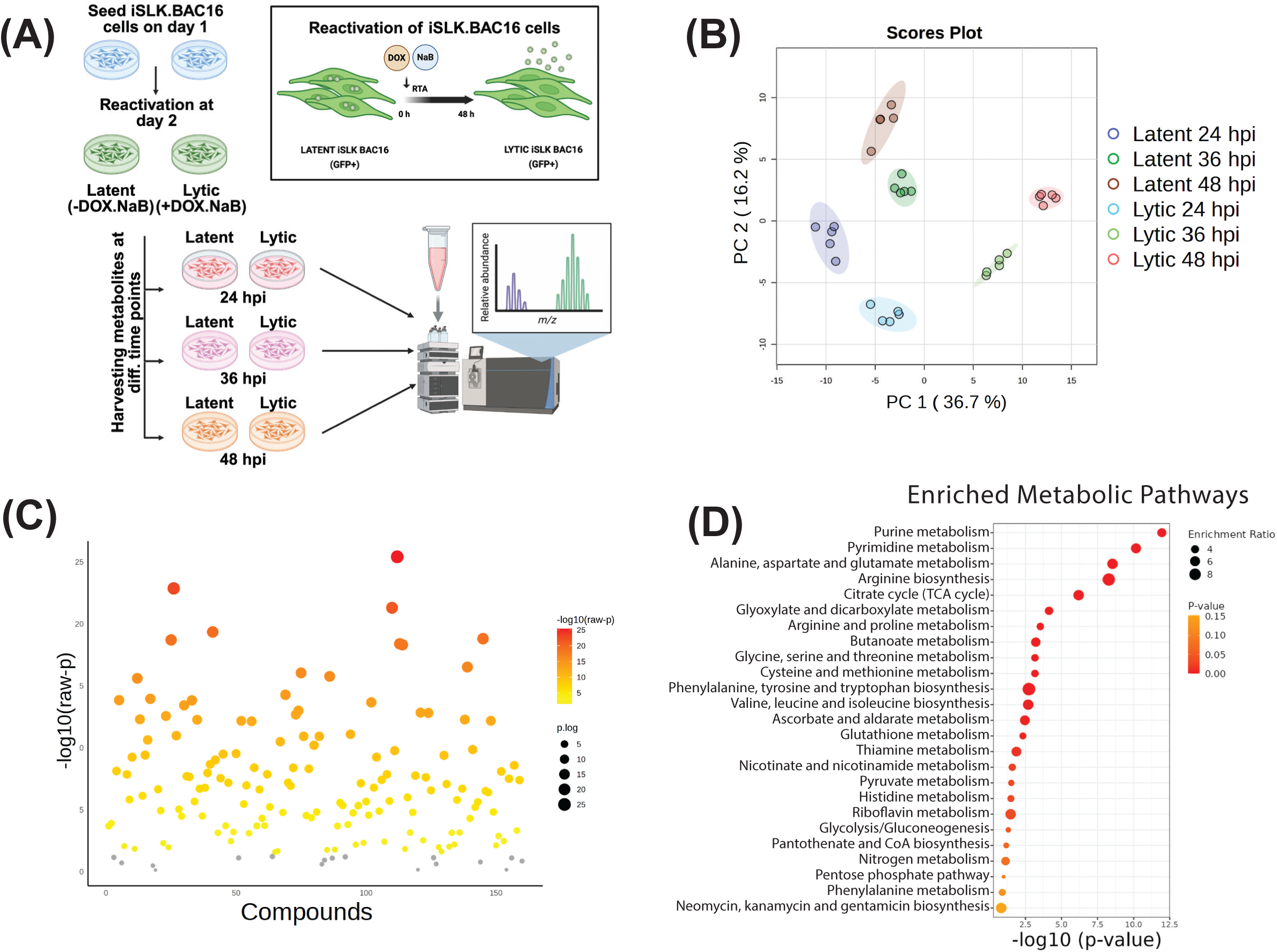
Targeted aqueous metabolomics of lytic vs latent KSHV-infected iSLK.BAC16 cells. (A) time-course experimental setup for a targeted aqueous metabolomics of lytic vs latent KSHV-infected iSLK.BAC16 cells using LC-MS technique (B) PCA analysis depicting the separation of the samples (C) The scatter plot illustrates the global pattern of significant changes in metabolites across samples. Statistical significance was assessed using one-way ANOVA with Tukey’s HSD post hoc test. FDR adjusted p-value (q-value) < 0.05 was considered significant. Color intensity represents ANOVA p-values, with darker shades indicating greater statistical significance while grey dots represent non-significant value (The full ANOVA statistics are provided in Supplemental Table 2) (D) Pathway enrichment analysis done on all the detected metabolites in the sample set, with purine and pyrimidine metabolic pathways being the most dysregulated. The size of the dots represents the enrichment ratio, which means the number of metabolite hits for that pathway. While the color of the dot represents the p-value, depicting the significance of that pathway being enriched. Significantly enriched pathways were identified using the “Pathway Enrichment Analysis” module based on the KEGG human reference library. Pathways are ranked by –log10(p-value), with larger values indicating greater statistical significance. Circle size reflects pathway impact based on topology analysis, and color indicates the degree of enrichment (from low to high).

### KSHV lytic infection alters amino acid, central carbon, and nucleotide metabolism

Upon deeper analysis of global metabolomics data from latent versus lytic KSHV infection, we find that the majority of metabolic pathways are altered during lytic infection. KSHV lytic infection alters amino acid metabolism, glycolysis, lactate production, glutaminolysis, fatty acid metabolism, and nucleotide metabolism. There are more amino acid monomers in lytic infection at different hours post-induction (hpi), i.e., 24, 36, and 48 hpi (**Figure 2A**). At 48 hpi, we measured an increase in amino acid monomers in lytic samples compared to latent. The essential amino acids histidine, isoleucine, leucine, methionine, phenylalanine, threonine, tryptophan, and valine have higher abundances in lytic samples at 36 and 48 hours post-induction. The metabolites associated with central carbon pathways are highly abundant in lytic KSHV samples when compared to latent **(Figure 2B).** Additionally, we measured an increase in glycolytic metabolites, including glucose-6-phosphate (G6P), phosphoenolpyruvate (PEP), and pyruvate, and energy-related metabolites like NAD+, NADPH, and FAD during KSHV lytic infection (**Figure 2B**). In lytic KSHV samples, we also measure an increase in TCA metabolites, mostly at 24 and 36 hpi, which are involved in reactions producing energy-rich molecules (NAD+ and FAD), such as isocitrate, alpha-ketoglutarate, fumarate, and malate. Furthermore, metabolites of nucleotide metabolism demonstrated differential abundance across the time course **(Figure 2C)**. At both 24 hpi and 48 hpi, we see a trend in the increase of nucleotide metabolites, precursors of DNA and RNA: dCTP, dATP, dTTP, UTP, CTP, ATP, and GTP. The nucleotide metabolites show a cumulative increase at both 24 and 48 hpi, suggesting an increased nucleotide metabolism during lytic infection. The UDP sugar pool essential for gluconeogenesis or glycosylation of proteins (UDP-glucose, UDP-glucuronate, and UDP-N-acetyl-glucosamine) was increased at 36 and 48 hpi during lytic infection(32) (**Figure 2C**).

**Figure 2.**
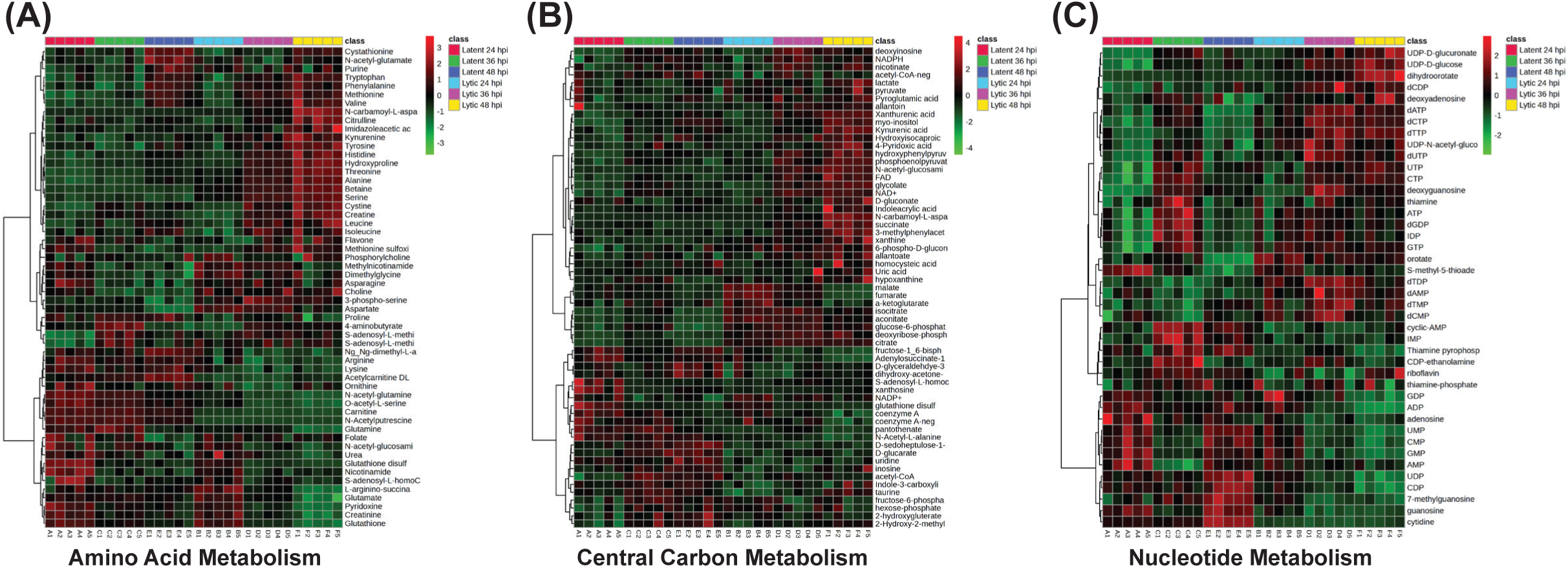
KSHV lytic infection alters key metabolic pathways. KSHV lytic infection induces metabolic alterations in host cells, including (A) amino acid, (B) carbohydrate, and (C) nucleotide metabolism. PCA plots show separation between samples. Metabolomics heat maps and PCA plots were generated using MetaboAnalyst 6.0, where the results were further normalized by sum and data were auto-scaled [(value-mean)/SD].

### Lytic KSHV infection increases DNA and RNA nucleotide biosynthesis

Deoxynucleotide triphosphates (dNTPs) are used to synthesize DNA (DNA replication), while nucleotide triphosphates (NTPs) are used in RNA synthesis (transcription) (33). DNA and RNA, being polynucleotide chains, are composed of purines-adenine and guanine; and pyrimidines-thymidine, cytidine, and uridine. In our LC-MS data, we report that mono- and di-phosphate deoxynucleotides (dNMPs and dNDPs) show a significant increase at 36 hpi, but their levels are reduced at 48 hpi compared to latent samples **(Figures 3A-B)**. However, the dNTPs (dATP, dCTP, and dTTP) show significant elevation at 36 hpi, and their levels remain elevated through 48 hpi when compared to earlier timepoints (**Figure 3C**). Interestingly, dNTPs remain elevated at 48 hpi, which could be due to the conversion of dNMPs and dNDPs into dNTPs to fulfil the need of the high DNA replication mechanism during lytic KSHV infection **(Figure 3C)**.

**Figure 3.**
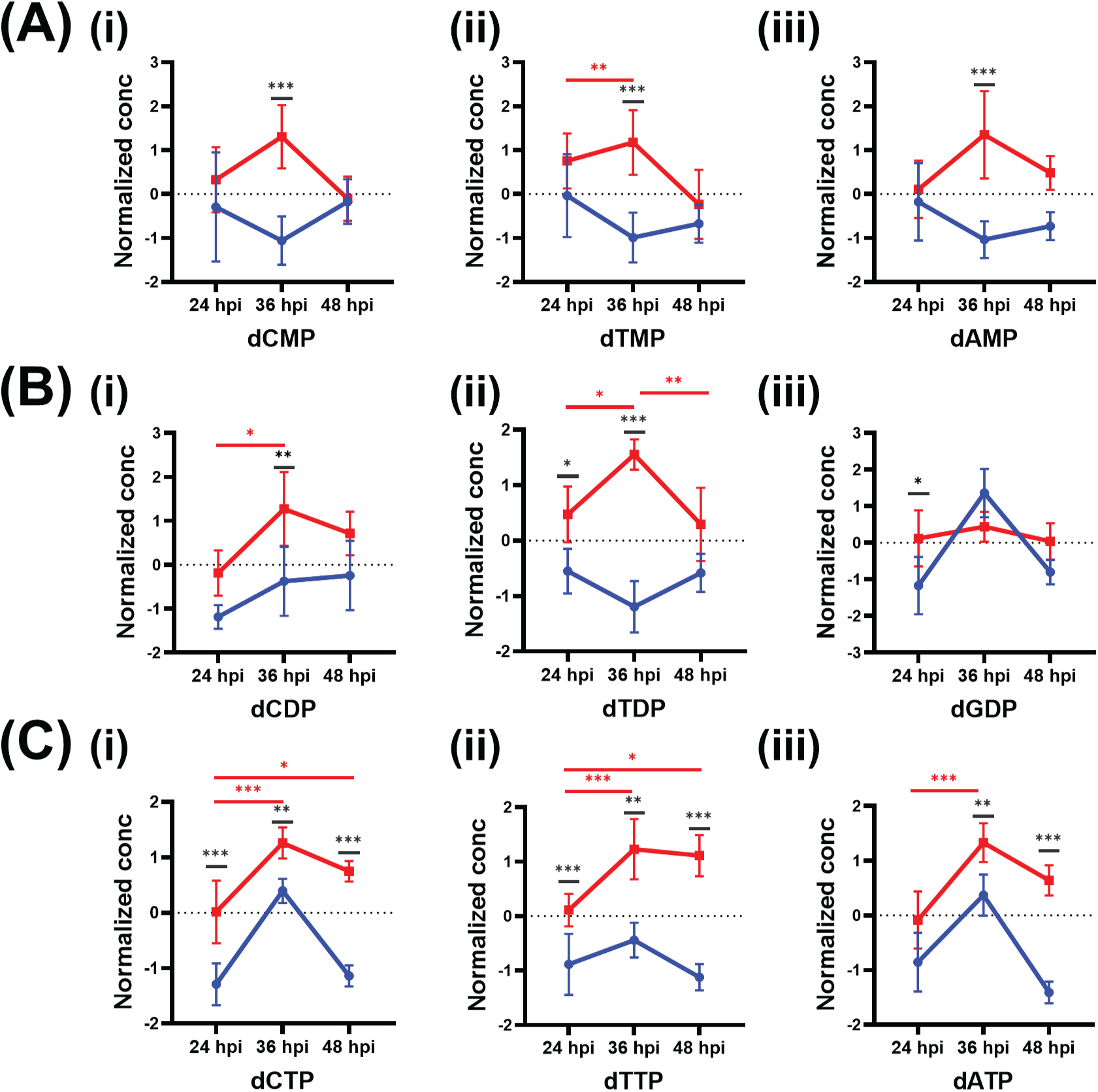
Deoxy-nucleotide metabolites are altered during lytic KSHV infection for maximal viral replication. iSLK.BAC16 cells, treated with or without Dox/NaB for reactivation, were harvested at 24, 36, and 48 hours post-induction (hpi). Deoxynucleotide phosphate species, (A) dCMP, dTMP, and dAMP; (B) dCDP, dTDP, and dGDP; were significantly abundant at 36 hpi when compared to latent infection, while (C) dCTP, dTTP, and dATP were significantly elevated at both 36 and 48 hpi compared to latent infection. Red lines represent lytic KSHV-infected samples, and blue lines represent latent KSHV infection. ***, P ≤ 0.001; **, P ≤ 0.01; * ≤ 0.05; n = 5

Next, we see mono- and di-nucleotide phosphates (NMPs and NDPs) are significantly low at 36 and 48 hpi with respect to 24 hpi **(Figure 4A-B)**. Triphosphate pyrimidines (UTP and CTP) and purines (ATP and GTP) are incorporated during RNA production. NTPs, especially pyrimidines, are seen to be significantly elevated in lytic infection at 48 hpi when compared to 24 hpi, as shown in the box plots (**Figure 4C**). Whereas, at 48 hpi, during lytic infection of iSLK.BAC16 cells, NMPs (CMP and CDP) **(Figure 4A)** and NDPs (CDP and GDP) **(Figure 4B)** are significantly low in abundance, and NTPs (CTP and GTP) **(Figure 4C)** are significantly high when compared to latent infection. This elevation in NTPs suggests that the KSHV virus requires active nucleotide metabolites (triphosphate species) for a high rate of DNA transcription and RNA production at 48 hpi.

**Figure 4.**
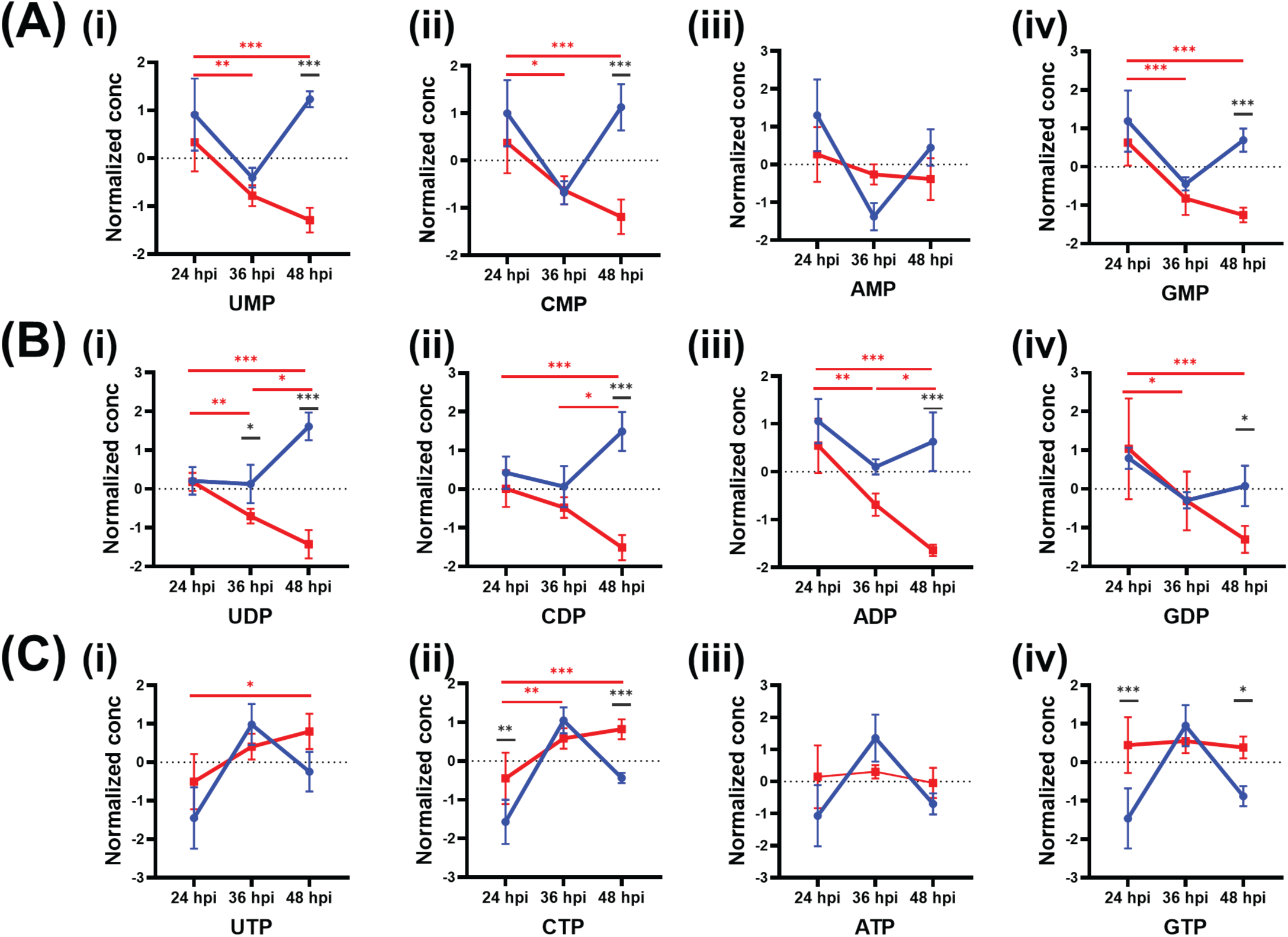
Nucleotide metabolites are altered during lytic KSHV infection for maximal virion production. iLSK.BAC16 cells, treated with or without Dox/NaB, were harvested at 24, 36, and 48 hours post infection. Nucleotide phosphate species, (A) UMP, CMP, AMP, GMP (B) UDP, CDP, ADP, GDP (C) UTP, CTP, ATP, GTP were significantly elevated at 48 hpi compared to latent infection. Red lines represent lytic KSHV-infected samples, and blue lines represent latent KSHV infection. ***, P ≤ 0.001; **, P ≤ 0.01; *, P ≤ 0.05; n=5

Apart from being the key unit of RNA polymer, ATP is the vital energy currency of the cells for all biosynthetic pathways (34). GTP is converted into ATP during times when high cellular energy is required. The box plots of ATP and GTP show an elevation at all time points (24, 36, and 48 hpi) during lytic infection when compared to latent infection **(Figure 4C)**.

### Nucleotide metabolism is required for maximal virion production during lytic gammaherpesvirus infection

To understand the requirement of nucleotide metabolism during KSHV lytic infection, we used an FDA-approved methotrexate (MTX), an anti-metabolite drug. MTX, a folate antagonist, competitively inhibits the host enzyme dihydrofolate reductase (DHFR). DHFR is required to convert folic acid to its active form, dihydrofolate (DHF, FH_2_), and tetrahydrofolate (THF, FH_4_). MTX treatment creates depletion of THF cofactors, including 5-methyl-THF, and thereby blocks several folate-related metabolic processes, including inhibition of nucleotide metabolism **(Figure 5A**). THF cofactor depletion in the host cells affects *de novo* synthesis of pyrimidine, deoxythymidine, required for DNA synthesis, and *de novo* synthesis of purine, guanosine, and adenosine, required for both DNA replication and transcription. Administering folinic acid (FA, 5-formylTHF) replenishes THF cofactors by bypassing rate-limiting metabolic enzymes rapidly converting into 5,10-MTHF, 5-MTHF, and THF (35). To test if elevated nucleotide metabolism is required for infectious KSHV virion production, we treated iSLK.BAC16 cells with 125nM MTX, which displayed minimal cytotoxicity, while supplementing the cells with folinic acid in the presence of MTX during lytic infection. Cell-free supernatants were collected after 48 hpi and titered on iSLK cells to determine viral production. The infected iSLK cells were washed twice with PBS and trypsinized, and the cells were resuspended in FACS buffer. After fixing with 4% formaldehyde (PFA), samples were analyze for FITC expression by flow cytometry. For each sample, 10,000 cellular events were recorded. For control samples treated with only DOX and Nab, ∼4% of cells expressed FITC/GFP **(Figure 5B)**. However, in lytic samples treated with MTX, there was a significant decrease in FITC-expressing cells, suggesting that there was a significant decrease in viral load after MTX treatment during lytic infection. By adding the downstream product of the DHFR enzymatic step, folinic acid, during MTX treatment in lytic samples, we see a significant increase in FITC-expressing cells to ∼14%. This data confirms that nucleotide metabolism is required for infectious KSHV virion production **(Figure 5C)**. However, the infectious virion particles increased when folinic acid was added alongside MTX treatment during lytic infection (**Figure 5C**). Titer images of supernatant from iSLK.BAC16 on iSLK shows the representative KSHV virion infection from each sample condition (**Figure 5D**). GFP+ cells indicate infected cells that have taken up the infectious KSHV virions. There is a notable decrease in GFP expression in MTX-treated samples; however, when folinic acid is supplemented, we see an increase in GFP+ cells, which represent more KSHV infection **(Figure 5E)**.

**Figure 5.**
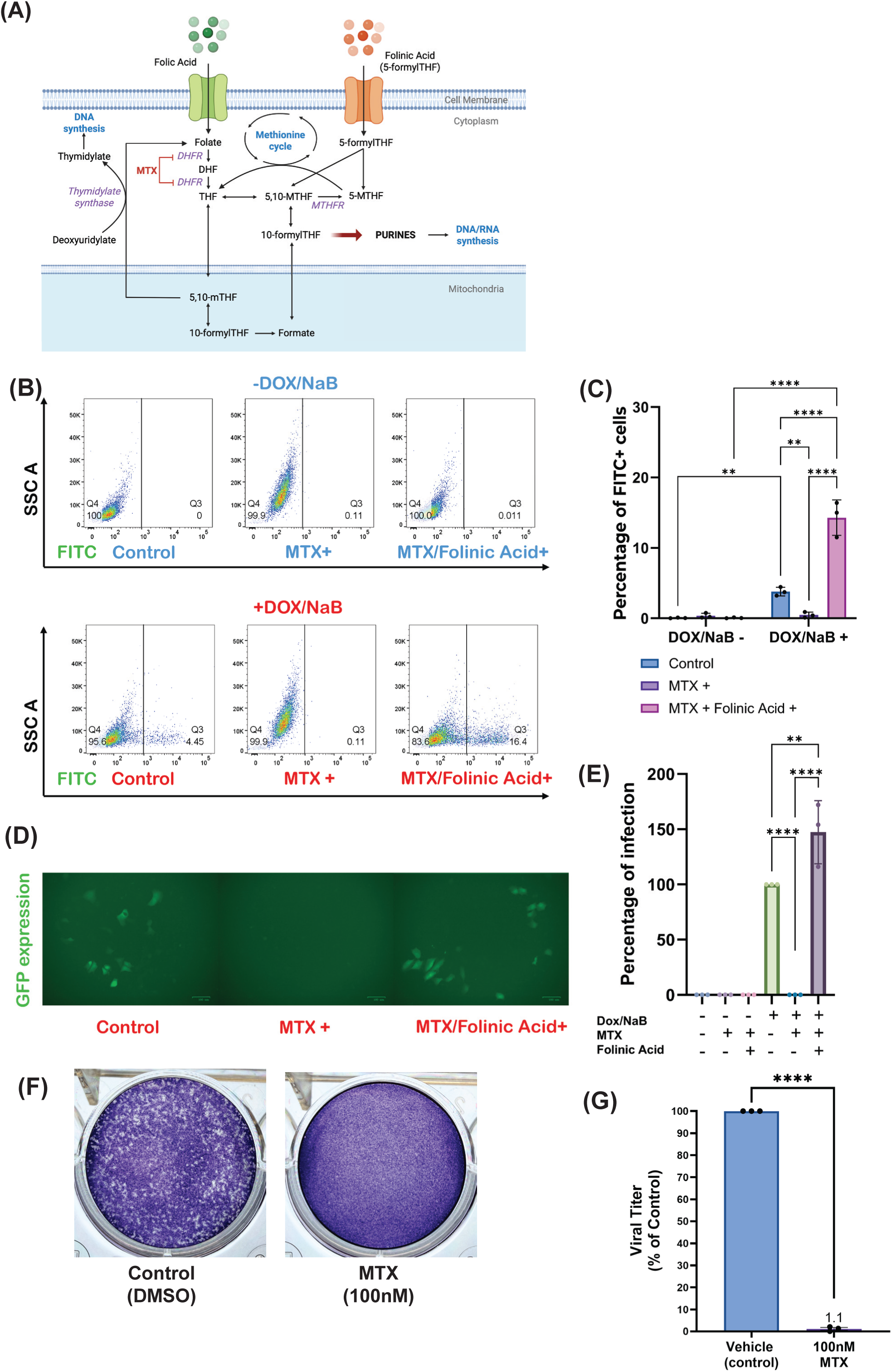
Methotrexate inhibition of nucleotide metabolism decreases infectious KSHV production. (A) Mechanism of action of methotrexate (MTX) and folinic acid (FA) during folate metabolism in host cell (B) iSLK.BAC16 cells were reactivated with 1μg/mL Dox and 1 mM NaB for 48 h. The cells were treated with methotrexate in the presence and absence of folinic acid. Reactivation rates increase in samples where the effects of methotrexate were rescued by folinic acid, as measured by the percentage of FITC+ cells. Flow data were analyzed using FlowJo software with latent samples as controls, with 10,000 incidents measured. The percentage of FITC+ cells decreased from ∼4% in the control to 0.4% in MTX-treated lytic cells. The percentage of FITC+ cells increased in FA-rescued MTX-treated lytic samples by ∼10% compared to the control. (C) Representative flow plots of GFP expression/FITC+ signal in iSLK.BAC16 cells (D) Images of lytic samples treated with MTX showed decreased extracellular viral titers, while FA treatment showed more GFP+ cells (E) Quantification of infectious KSHV extracellular viral titer images showed a significant decrease during MTX treatment and a significant increase post-FA treatment of MTX lytic samples. (F - G) NIH 3T3 cells were MHV-68 infected (MOI = 0.1) and treated with vehicle (DMSO control) or 100 nM MTX for 48 hours in three independent experiments. Viral titer was quantified from extracellular supernatants via plaque assays. F) Representative MHV-68 plaque assay images from vehicle (control) or 100 nM MTX-treated cells. G) Quantification of MHV-68 extracellular viral titer as a percentage of the control (vehicle treatment). ****, P ≤ 0.0001; ***, P ≤ 0.001; **, P ≤ 0.01; *, P ≤ 0.05; n=3

As previously noted, metabolomics analysis revealed lytic MHV-68 infection increases nucleotide metabolism (25). To test if MTX treatment also reduced lytic infectious virus production during mouse gammaherpesvirus infection, we mock- or MHV-68-infected (MOI = 0.1) NIH 3T3 cells and then treated them with DMSO (vehicle control) or 100 nM MTX for 48 hrs. While 100 nM MTX treatment reduced cell proliferation compared to vehicle (control), 100 nM MTX treatment resulted in a 99% cell viability in mock- and MHV-68 infected NIH 3T3 cells (**Supplemental Table 3**). Quantitative plaque assay analysis revealed that 100nM MTX treatment of MHV-68-infected cells decreased infectious virus production by 98.9% compared to control-treated cells (Fig. 5F - G). After normalization of the data of viral titer to live cell number, our data showed a ∼98.5-fold decrease in virus production compared to control-treated cells. Overall, these data suggest the induction of nucleotide metabolism by both human (KSHV) and mouse (MHV-68) gammaherpesvirus are necessary for virion production.

### Nucleotide metabolism is required for late lytic KSHV gene expression

To determine how MTX affects lytic infection, the iSLK.BAC16 cells were treated with and without 125 nM MTX, which displayed minimal cytotoxicity in the presence or absence of Dox/NaB to induce the lytic phase **(Supplemental Figure 1)**. The cells were reactivated at the same time as MTX treatment to make sure nucleotide synthesis was inhibited, and there was no further *de novo* production of nucleotides during reactivation. FA was supplemented in the presence or absence of MTX during reactivation. After 48 hpi, the supernatant was collected and saved to perform titer experiments. RNA was extracted from cells, and viral gene expression was quantified using RT-qPCR. The relative mRNA expression level of latent gene LANA showed a significant increase after MTX treatment, and FA supplementation decreased LANA mRNA expression **(Figure 6A)**. Next, we analyzed the mRNA expressions of the early lytic gene, ORF59, and a late lytic gene, K8.1. ORF59 mRNA expression significantly increased after MTX treatment, while FA treatment showed varied expression across biological replicates, exhibiting no significant change (**Figure 6B**). Interestingly, we measured a significant decrease of about 30% in K8.1 relative mRNA levels (**Figure 6C**). FA treatment significantly reduced MTX anti-viral effects, and there was an increased expression of 50% of K8.1 compared to the control when both MTX and folinic acid were administered to the cells during lytic infection (**Figure 6C**). These findings suggest that host nucleotide metabolism is required for KSHV lytic infection. The drug inhibition of nucleotide metabolism as well as its rescue using a substrate metabolite show that KSHV lytic infection is dependent on the host metabolic pathways in order to produce infectious KSHV virions.

**Figure 6.**
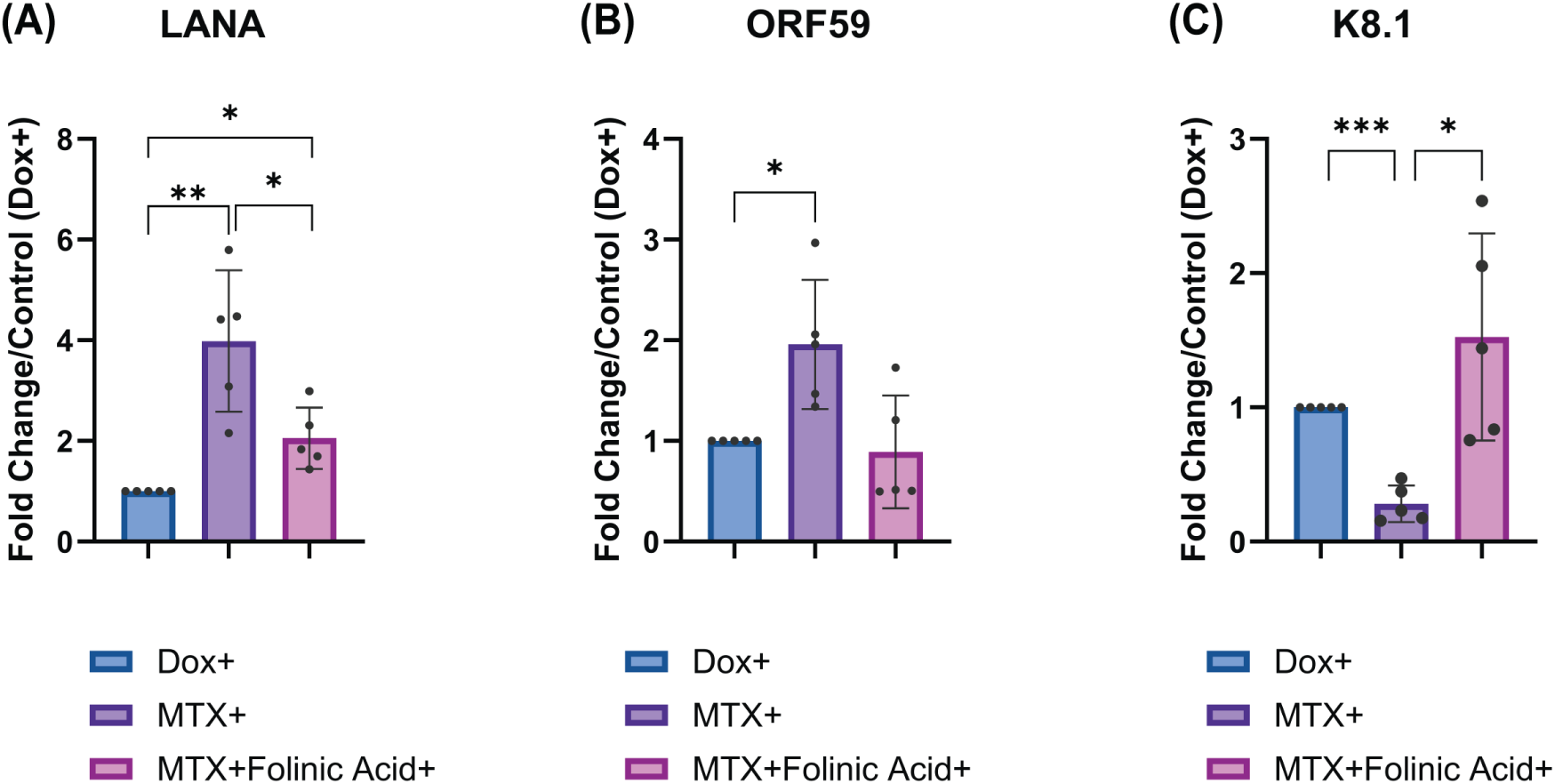
Methotrexate inhibition of nucleotide metabolism decreases KSHV late lytic gene expression. iSLK.BAC16 cells were induced in the presence or absence of 125 nM MTX, and FA was supplemented to the MTX samples. Cells were harvested at 48 hpi. RNA was extracted, and qRT-PCR was performed on cDNA for (A) latent KSHV transcript (LANA), (B) early KSHV transcript (ORF59), and (C) late KSHV transcript (K8.1). Relative mRNA expression for each viral transcript was normalized to host actin as the housekeeping gene and compared to that of untreated induced samples. Error bars are representative of standard errors from 5 biological replicate experiments. ***, P ≤ 0.001; **, P ≤ 0.01; *, P ≤ 0.05; n=5

## Discussion

In this study, we demonstrate that nucleotide metabolism is elevated and required by KSHV lytic infection for maximal virion production and thus represents a potential therapeutic target for treating KSHV infection. Alterations in the global host metabolome have been previously studied in the presence of other viral infections like HCMV, vaccinia virus (VACV), HCV, EBV, and HSV-1, as well as KSHV latent infection (13, 16–19, 22, 36). Our PCA analysis shows a clear separation between latent and lytic samples, which indicates a unique metabolic profile of KSHV infection at 24, 36, and 48 hpi in both viral phases **(Figure 1B)**. In this paper, we have defined the altered metabolic pathways in KSHV lytic infection compared to latent at various time points post lytic induction **(Figure 1C)**. Using LCMS targeted metabolomics on KSHV latent and KSHV lytic samples collected at several time points, we were able to perform data analysis and pathway enrichment analysis to determine significantly altered metabolic pathways **(Figure 1D)**. We found that the pathway altered the most in KSHV lytic samples was nucleotide metabolism. Other top pathways altered during KSHV lytic infection included amino acid metabolism, the citrate cycle, glutathione metabolism, arginine metabolism, and thiamine metabolism. Glutathione metabolism is required to maintain the redox homeostasis of the host cell. KSHV-encoded microRNAs regulate the expression of xCT transporter (SLC7A11) and protect the cells from reactive nitrogen species (RNS) to protect the cells from extreme redox environments and help KSHV persist in infection(37). Another study confirms that the KSHV ORF 59 lytic gene modulates histone 4 arginine methylation, which controls the compactness of the chromatin and induces lytic reactivation, thus altering the arginine metabolic pathway (38).

In our LCMS dataset, we see an increase in anabolic metabolites during KSHV lytic infection. An increase in the abundance of essential amino acids—histidine, isoleucine, leucine, methionine, phenylalanine, threonine, tryptophan, and valine—suggested the demand for protein synthesis to fulfill viral replication needs **(Figure 2A)**. A study done on HSV-1 demonstrated the requirement of essential amino acids for higher viral load (39). Among the essential amino acids in the LCMS data, only lysine was decreased in lytic KSHV infection. Lysine residues are the major sites for post-translational modification (PTM) of both viral and host proteins, which KSHV manipulates to dysregulate PTM and evade host immune response (40–42). Interestingly, we see higher glutamine presence at 24 and 36 hpi in latent samples. Previous groups have established that KSHV latent infection induces glutaminolysis and enhances glutamine uptake (43, 44) **(Figure 2A)**. Further analysis of the upregulated metabolic pathways showed an increase in the energy-related metabolites of NAD+, NADPH, and FAD during lytic KSHV infection and an increase in the metabolites that are involved in reactions with these energy metabolites, such as glucose-6-phosphate (G6P), phosphoenolpyruvate (PEP), and pyruvate in glycolysis, and isocitrate, alpha-ketoglutarate, fumarate, and malate in the TCA cycle (**Figure 2B**). Furthermore, an increase in lactate abundance implies that there is a Warburg effect occurring in KSHV lytic samples. The Warburg effect is defined as the cellular usage of glycolysis for energy production instead of oxidative phosphorylation, even in the presence of oxygen, in order to get rapid energy production and ensure viral replication(20). The role of the Warburg effect has been established in latent KSHV infection in previous studies (20). The increase of nucleotide-related metabolites at both 24 and 48 hpi suggests an increased requirement of nucleotide metabolism during KSHV lytic infection **(Figure 2C)**. As the KSHV replication cycle includes two phases of DNA replication, immediate early and early lytic gene replication and then genome replication, we hypothesize that an increase in nucleotides at 24 hpi is to compensate for early gene expressions. But a more pronounced difference in nucleotides at 48 hpi is due to genome replication. Since genome replication is also followed by transcription and translation of viral proteins, it further suggests that the increase in amino acids at 48 hpi is for viral protein formation **(Figure 2C)**. Our LC-MS data show significant increased abundances of these UDP-sugars at 36 and 48 hpi during lytic KSHV infection **(Figure 2C)**. All herpesviruses, including KSHV, have glycoproteins on the outermost lipid membrane for initiating binding of host cell receptors either during new infection or egress from the host cell (45–47). UDP sugars such as UDP-glucose, UDP-glucuronate, and UDP-N-acetyl-glucosamine serve as glycosylation substrates during post-translational modifications(48). KSHV encodes for eight glycoproteins, which are incorporated in the virion envelope: ORF8 (gB [glycoprotein B]), ORF28, ORF68, ORF22 (gH), ORF47 (gL), ORF39 (gM), ORF53 (gN), and gpK8.1(49, 50). These glycoprotein structures share homology with other herpesviruses or are unique to KSHV itself (51–53).

Finally, our data show that deoxynucleotides are significantly higher in abundance at 36 and 48 hpi during lytic infection compared to latent samples, suggesting that DNA replication occurs throughout lytic KSHV infection **(Figure 3)**. Upon reactivation, an organized cascade of gene expression is initiated with the immediate early and early lytic gene expression, viral genome replication, and late lytic gene expression(54). The host metabolomic machinery aids KSHV for viral DNA replication and gene expression(55). However, the increase in nucleotide phosphate species at 48 hpi suggests that there is more genome transcription required during the later phase of the viral lytic replication cycle **(Figure 4)**. We determined the requirement for nucleotide metabolism during lytic infection by demonstrating that inhibiting *de novo* nucleotide production indirectly with the chemical MTX reduced late lytic viral gene expression, K8.1, and decreased extracellular virion production. MTX, a folate antagonist, binds tightly (competitively) and inhibits the host enzyme dihydrofolate reductase (DHFR). DHFR is a key enzyme in folate metabolism, which recycles folic acid to THF (56). MTX treatment creates depletion of THF cofactors and thereby blocks several folate-related metabolic processes, one of which is inhibition of nucleotide metabolism (57). MTX has a higher affinity to DHFR than folic acid and hence is used at low cellular doses to decrease death by toxicity (30, 58). MTX inhibition of DHFR bypassed by the administration of folinic acid (FA, 5-formyl THF), which is a reduced stable form of folic acid, restores the folate derivatives and decreases the toxic effects of MTX (59). Notably, lytic KSHV encodes for viral DHFR (ORF2), which is catalytically more active and does not show inhibition by known inhibitors of the DHFR enzyme, including MTX (57, 60). During lytic KSHV infection, supplementing cells with FA during MTX inhibition, we see a significant increase in the relative mRNA expression of late lytic viral gene expression, K8.1, compared to control **(Figure 5)**. FA-supplemented cell-free supernatant titered on uninfected cells showed a significant increase in extracellular virion production. **(Figure 6)**.

In conclusion, our findings define and expand our understanding of the host metabolic profile during lytic gammaherpesvirus infection **(Figure 7)**. We demonstrate a unique metabolic profile of host cells undergoing lytic KSHV infection at 24, 36, and 48 hours post-induction. Gammaherpesviruses, including KSHV, MHV-68, and EBV, exploit host nucleotide metabolism to establish persistent viral infection and replication, leading to a diseased state (24, 25, 61). Reduction in viral titers by MTX-mediated nucleotide metabolic inhibition shows that there is a shared metabolic vulnerability among gammaherpesviruses. The findings of this study provide a mechanistic insight into how gammaherpesviruses exploit host metabolic pathways to sustain lytic replication.

**Figure 7.**
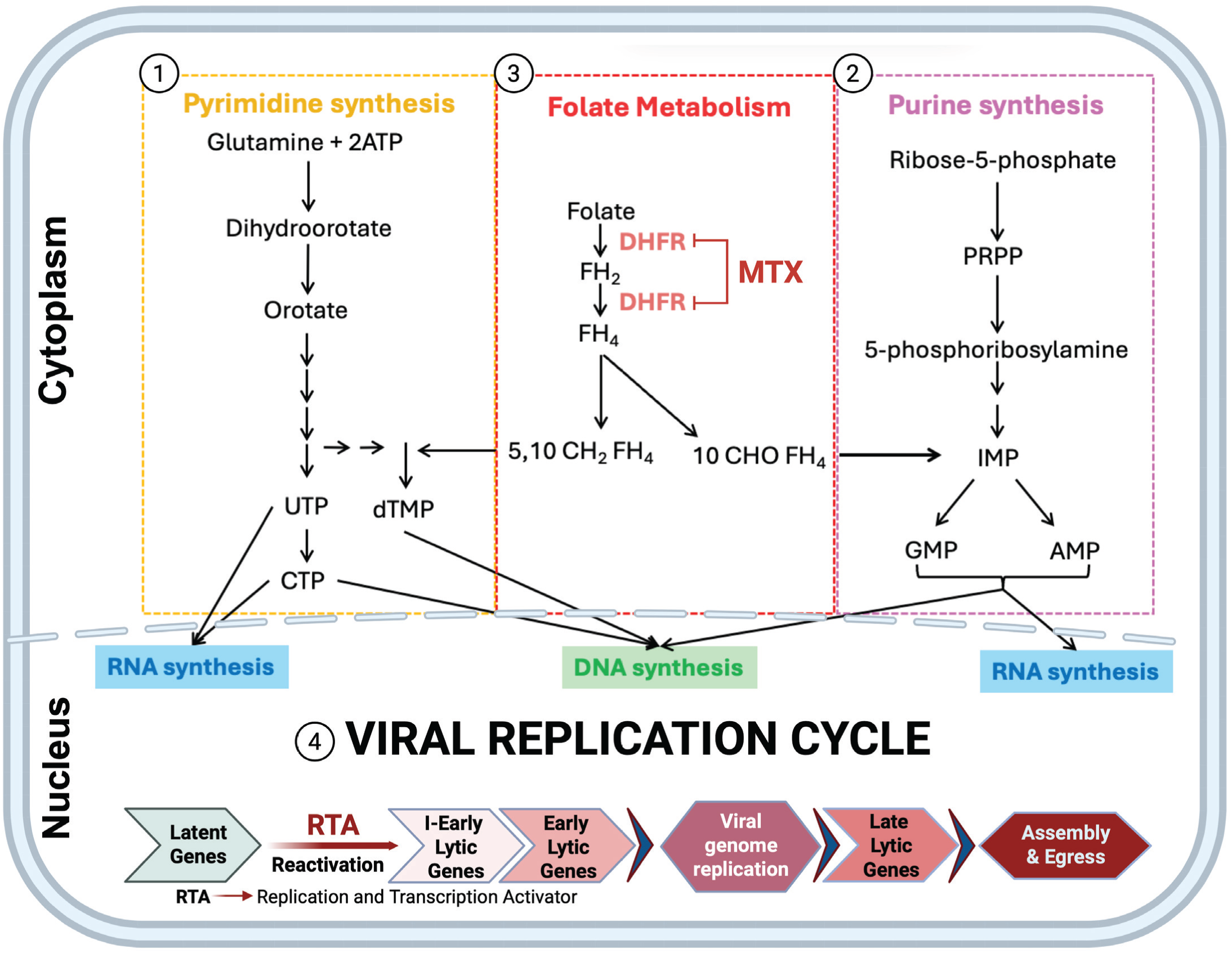
Summary model of host cell nucleotide metabolism changes during lytic gammaherpesvirus infection. Our time-course global metabolomics indicate that during KSHV lytic infection of iSLK.BAC16 cells, metabolic intermediates found in nucleotide metabolism are significantly elevated in lytic samples when compared to latent. In host cell, *de novo* nucleotide synthesis includes pyrimidine and purine synthesis which occurs in the host cytoplasm. This figure illustrates how de novo pyrimidine and purine biosynthesis pathways support viral replication following reactivation from latency. (1) Pyrimidine synthesis (left) generates UTP, CTP, and dTMP, while (2) purine synthesis (right) produces IMP-derived GMP and AMP; both pathways converge on nucleotide pools required for viral RNA and DNA synthesis. (3) Central folate metabolism (red box), which supplies one-carbon units for thymidylate and purine production, is shown as a key regulatory node catalysed by dihydrofolate reductase (DHFR) and inhibited by methotrexate (MTX). (4) In the nucleus (bottom), the viral replication cycle is depicted, beginning with RTA-mediated reactivation, followed by immediate-early, early, and late lytic gene expression, viral genome replication, and assembly/egress. Together, the schematic highlights that robust nucleotide biosynthesis is essential for efficient herpesvirus lytic replication.

**Supplemental Figure 1. Methotrexate exhibits minimal cytotoxic effects at 125 nM.** iSLK.BAC16 cells were treated with control (DMSO) and different concentrations of MTX: 50nM, 125nM, 250nM, and 375nM. After 48 hours, the cell viability was calculated as [(live cells/total cells)*100 = cell viability %] where: total cells = live + dead cells.

**Supplemental Table 1. Time course metabolomics data normalized to protein concentration.**

**Supplemental Table 2. ANOVA statistics**

**Supplemental Table 3. Methotrexate exhibits minimal cytotoxic effects at 100 nM in mock- and MHV-68 infected NIH3T3 cells.**

## Material and Methods

### Cells and Reagents

iSLK.BAC16 cells(26) or iSLK cells(62) were maintained at 37℃ in 5% CO₂ as monolayer cultures in Dulbecco’s modified Eagle medium (DMEM) (Corning, 10-013-CV) with L-Glutamine (Corning, 25030-081) and sodium pyruvate, additionally supplemented with 10% fetal bovine serum (FBS) (Corning, 35-011-CV) and 1% L-Glutamine. iSLK.BAC16 cell medium was selected with puromycin (10 mg/ml), G418 (95 mg/ml), and hygromycin B (Corning, 30-240-CR; 50 mg/ml). iSLK cell medium was supplemented with puromycin (10 mg/ml) and G418 (95 mg/ml). All KSHV lytic experiments were carried out in DMEM with 10% FBS and 1% L-Glutamine, without selective agents.

For MHV-68 studies, NIH 3T3 cells (ATCC no. CRL-1658) or Vero cells (ATCC no. CCL-81) were cultured at 37°C and 5% CO_2_ in DMEM containing high glucose, L-glutamine, sodium pyruvate (Genesee no. 25-500), 1% penicillin streptomycin (Genesee no. 25-512). Additionally, NIH 3T3 cells were supplemented with 10% newborn calf serum (Fisher #16010159) and Vero cells were supplemented with 10% fetal bovine serum (Genesee no. 25-550).

### Virus

The iSLK.BAC16 cell line stably maintains the KSHV genome. The cells are latently infected with recombinant KSHV.BAC16, which encodes constitutive expression of EGFP. These cells also encode the viral Replication and Transcription Activator (RTA) transgene, which is doxycycline (1ug/mL) (Dox) and sodium butyrate (1ug/mL) (NaB) inducible; this triggers the transition in iSLK.BAC16 from latent to lytic infection. iSLK cells do not contain the KSHV genome and, therefore, are used as a control in our experiments.

### Drug treatment on KSHV infection

Methotrexate (MTX) (ThermoScientific Chemicals, J63075.MC) and folinic acid were prepared in DMSO and were used at a final concentration of 125 nM and 2 uM in DMEM, respectively. iSLK.BAC16 cells were seeded on day 1 and reactivated (Dox and NaB) the next day. MTX and folinic acid were added at the time of reactivation. At 48 hours post-treatment, a cell-free supernatant was collected and saved to perform virus titer experiments. The cells were harvested for qPCR analysis.

### MHV-68 infection, MTX treatment, viral titer, and cell counts

MTX (Sigma no. 1414003) was dissolved in dimethyl sulfoxide (DMSO) and stored at −80°C until use. NIH 3T3 cells were seeded at a density of 760,000 cells in 6 cm dishes. MHV-68 viral stocks (ATCC no. VR-1465) were propagated via infection of NIH 3T3 cells or Vero cells at an MOI of 0.01-0.05 as previously described (25). Approximately 4 hours post seeding, NIH 3T3 cells were mock- or MHV-68 infected (MOI = 0.1) in serum free DMEM for 2 hours and rocked every 20 minutes. After infection, the cell media was aspirated, replaced with 3mL fresh complete DMEM, and either vehicle (DMSO control) or 100 nM MTX. At 48 hpi, the viral supernatant was cleared by centrifugation (1,000 rpm) and frozen at −80°C for future plaque assay analysis. The cell pellets after centrifugation were combined with trypsinized cells from matching samples. Each sample 1) Mock (DMSO control), 2) Mock (100 nM MTX), 3) MHV-68 (DMSO control), and 4) MHV-68 (100 nM MTX) was centrifuged for 125 x g for 5 min. The media was aspirated and the cell pellet was resuspended in 500 µL complete DMEM. Total cells, live cells, and dead cell numbers were determined by a trypan blue exclusion assay using the automated Biorad T-20 cell counter (Biorad no. 1450102). Viral titers were quantified through traditional viral plaque assays and crystal violet staining as previously described (25). Plaque forming units per mL (pfu/mL) were calculated from duplicate wells using the equation: (dilution factor) x (1 / 0.2mL) x (average pfu) = pfu/Ml

### Quantitative reverse transcription-PCR (qRT-PCR)

All treated cells were washed once with Phosphate-Buffered Saline (PBS) (Corning, 21-040-CV), and RNA was extracted following the manufacturer’s manual using the PureLink™ RNA Mini Kit (ThermoFisher Scientific, 12183025). A two-step quantitative real-time reverse transcription PCR (RT-qPCR, Applied Biosystems) was used to measure mRNA expression levels of host metabolic genes and KSHV viral latent and lytic genes. Relative changes in gene expression were normalized using the delta threshold cycle (ΔCt) method to the abundance of host actin mRNA (housekeeping gene). (ΔCt) values were then used to determine ΔΔCt when compared to the dox-treated control group (iSLK.BAC16 cells treated with Dox and NaB).

### Primers

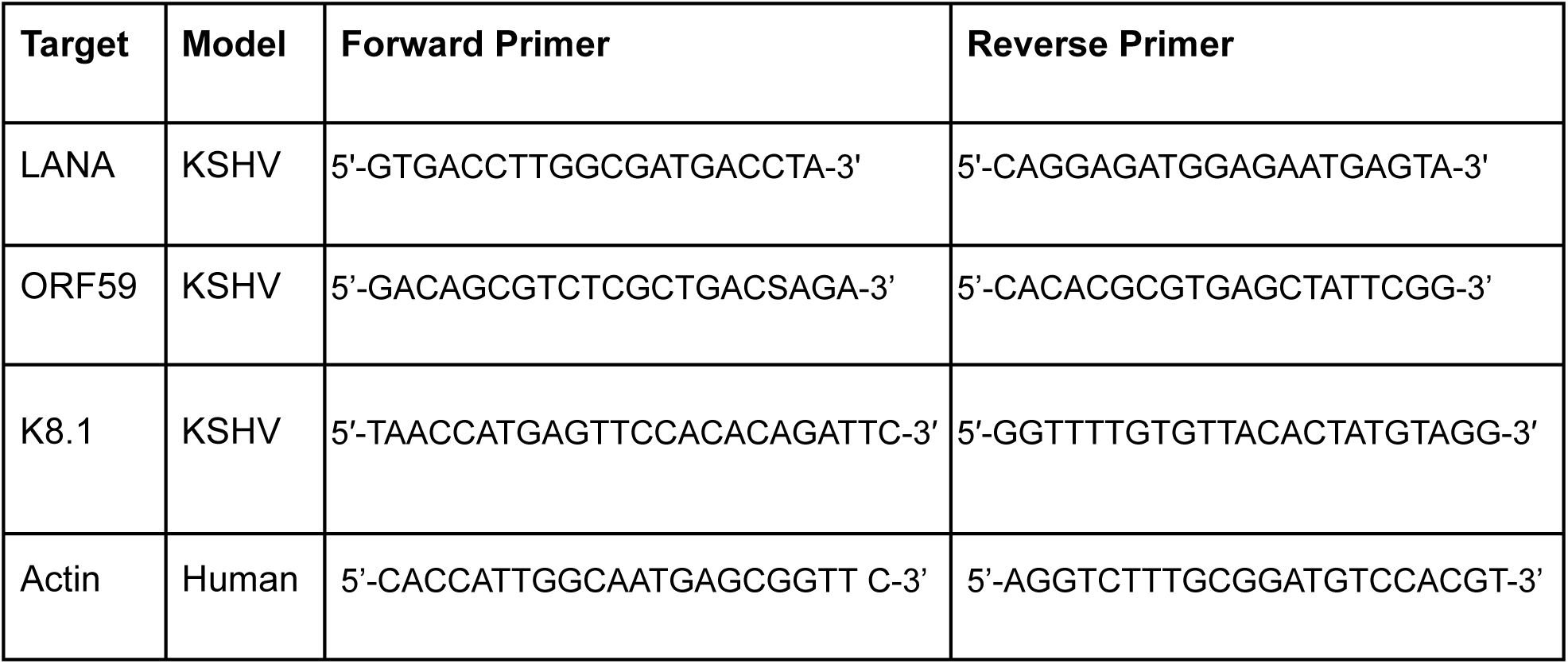

### Spinoculation Infection

All iSLK.BAC16 cell supernatants were collected post-experimental treatment (specifics described in the results section) and subjected to centrifugation for clearing debris at 300 X g for 10 minutes. iSLK cells were seeded in 12-well plates at 70% confluency 24 hours prior to induction. To titer, 500 µL of supernatant from treated iSLK.BAC16 cells, supplemented with polybrene (10 µg/mL), were added to iSLK cells.

Infection was enhanced by spinoculation at 300 X g, 37°C for 1 h, and then incubated for 4 hr at 37°C with interval rocking plates every 30 mins. Infectious supernatant media was replaced with fresh iSLK medium after 4 hours. Titer plates were imaged 48 hours post-infection at 10X magnification on the Zoe microscope (BIORAD more info) for GFP+ cells. GFP-positive cells were quantified using ImageJ, and a t-test was run between samples and control (Dox+). The significant p-values are denoted as follows: p value > 0.001 (**), p value < 0.0001(****)

### Flow cytometry

iSLK cells were seeded for spinoculation as described above. Forty-eight hours post-titer infection, cells were washed once with PBS, trypsinized, collected, and pelleted by centrifugation at 300 X g for 5 minutes. The supernatant was discarded, and the cell pellet was resuspended in 200 µL of FACS Buffer (2% FBS in PBS). The cells are fixed in 4% paraformaldehyde for 5 minutes, washed with 2 mL PBS, and centrifuged for 5 min at 300 X g. The supernatant was removed, and the pellet was resuspended in a 500 μL FACS buffer. For flow cytometry analysis, the samples were immediately analyzed using a flow cytometer equipped with filters suitable for detecting FITC. A BD Fortessa SORP cytometer was used for FACS data collection. During data collection, a minimum of 10,000 cells/events per sample were recorded to ensure statistical significance. In the data analysis phase, the acquired data are processed using FlowJo. The population was selected firstly for live cells and then for singlets. The singlet cells were then analyzed for FITC to determine the number of KSHV-infected cells (GFP+).

### Metabolite Extraction

iSLK.BAC16 cells were seeded in 60-mm dishes with 5 biological replicates per sample and harvested across a time course post reactivation (24, 36, and 48 hours). Cells were harvested for metabolomics, and supernatant was saved for titering. Briefly, cells were washed with chilled PBS three times on dry ice and at −80°C. Next, the metabolite extraction buffer, 80:20 methanol:water (v/v), was immediately added to quench cellular metabolism. The cells were scraped on dry ice, collected in microcentrifuge tubes, and centrifuged at a high speed (14,000 rpm) for 15 mins at 4°C. The pellets were saved for protein quantification, and the supernatant with metabolites was centrifuged twice at a high speed (14,000 rpm) for 15 mins at 4°C. Finally, the metabolite-containing solution was transferred to HPLC vials and transported to the UT Southwestern Medical Center Metabolomics Core Facility. Samples were stored at −80°C until processed on a mass spectrometer. The core facility provided the metabolic profiling of the samples using targeted metabolomics.

### LC-MS/MS instrumentation system

Mass spectrometric analyses were performed on a Sciex “QTRAP 6500+” mass spectrometer equipped with an ESI ion spray source. The ESI source was used in both positive and negative ion modes. The ion spray needle voltages used for MRM positive and negative polarity modes were set at 4800 V and −4500 V, respectively. The mass spectrometer was coupled to Shimadzu HPLC (Nexera X2 LC-30AD). The system is under control by Analyst 1.7.1 software.

### Chromatography Conditions

Chromatography was performed under HILIC conditions using a SeQuant® ZIC®-pHILIC 5 μm polymeric 150 × 2.1 mm PEEK-coated HPLC column (MilliporeSigma, USA). The column temperature, sample injection volume, and flow rate were set to 45°C, 5 μL, and 0.15 mL/min, respectively. The HPLC conditions were as follows: Solvent A: 20 mM ammonium carbonate including 0.1% ammonium hydroxide and 5 μM medronic acid. Solvent B: Acetonitrile. The gradient condition was 0 min: 80% B, 20 min: 20% B, 20.5 min: 80% B, 34 min: 80% B. Total run time: 34 mins. Data was processed by SCIEX MultiQuant 3.0.3 software with relative quantification based on the peak area of each metabolite.

### Metabolomics data analysis

The concentration of each metabolite (M) was divided by total protein (P) to normalize to external control as follows: C = M/P, where C is the normalized concentration of the metabolites. Metabolomics heat maps and PCA plots were generated using MetaboAnalyst 6.0, where the results were further normalized by sum and data were auto-scaled [(value-mean)/SD]. Box plots for nucleotide metabolites were generated using normalized concentration (C), and the t-test was run between latent and lytic samples as well as between different time points in latent and lytic samples. The significance is denoted as follows: p value > 0.05 (ns), p value > 0.01 (*), p value > 0.001 (**), p value < 0.001(***)

### Statistical Analysis

Figures and statistical analyses were generated using GraphPad Prism 8.0.1 software. Experiments were performed in at least three biological replicates. Data are presented as the mean ± SEM. KSHV experimental conditions were compared to control conditions using ANOVA. MHV-68 viral titer was analyzed using a paired Student’s (two-tailed) t test. Significant p-values are noted in each figure (* ≤ 0.05; ** ≤ 0.01; *** ≤ 0.001), and “ns” denotes a non-significant change.

## Acknowledgments

E.L.S. and F.H. were supported by the National Institutes of Health (NIGMS) 1R35GM160182. We would also like to thank the members of the Sanchez lab for their insightful discussion and input throughout the study. T.D. and Y.A.F were supported by the National Institutes of Health (NIH) 1R15AI185858 and the Seattle Pacific University Faculty Scholarship Grant. The Delgado lab would like to thank the 2024 Seattle Pacific University Medical Virology course undergraduate students for their preliminary findings showing MTX has MHV-68 antiviral activity.

## References

1. Cesarman E, Damania B, Krown SE, Martin J, Bower M, Whitby D. 2019. Kaposi sarcoma. Nat Rev Dis Primers 5:9.

2. Ganem D. 1997. KSHV and Kaposi’s sarcoma: the end of the beginning? Cell 91:157–160.

3. Dupin N, Fisher C, Kellam P, Ariad S, Tulliez M, Franck N, van Marck E, Salmon D, Gorin I, Escande JP, Weiss RA, Alitalo K, Boshoff C. 1999. Distribution of human herpesvirus-8 latently infected cells in Kaposi’s sarcoma, multicentric Castleman’s disease, and primary effusion lymphoma. Proc Natl Acad Sci U S A 96:4546–4551.

4. Ariyoshi K, Schim van der Loeff M, Cook P, Whitby D, Corrah T, Jaffar S, Cham F, Sabally S, O’Donovan D, Weiss RA, Schulz TF, Whittle H. 1998. Kaposi’s sarcoma in the Gambia, West Africa is less frequent in human immunodeficiency virus type 2 than in human immunodeficiency virus type 1 infection despite a high prevalence of human herpesvirus 8. J Hum Virol 1:193–199.

5. Renne R, Lagunoff M, Zhong W, Ganem D. 1996. The size and conformation of Kaposi’s sarcoma-associated herpesvirus (human herpesvirus 8) DNA in infected cells and virions. J Virol 70:8151–8154.

6. Trus BL, Heymann JB, Nealon K, Cheng N, Newcomb WW, Brown JC, Kedes DH, Steven AC. 2001. Capsid structure of Kaposi’s sarcoma-associated herpesvirus, a gammaherpesvirus, compared to those of an alphaherpesvirus, herpes simplex virus type 1, and a betaherpesvirus, cytomegalovirus. J Virol 75:2879–2890.

7. Purushothaman P, Dabral P, Gupta N, Sarkar R, Verma SC. 2016. KSHV genome replication and maintenance. Front Microbiol 7:54.

8. Ciufo DM, Cannon JS, Poole LJ, Wu FY, Murray P, Ambinder RF, Hayward GS. 2001. Spindle cell conversion by Kaposi’s sarcoma-associated herpesvirus: formation of colonies and plaques with mixed lytic and latent gene expression in infected primary dermal microvascular endothelial cell cultures. J Virol 75:5614–5626.

9. Liu X, Zhu C, Wang Y, Wei F, Cai Q. 2021. KSHV reprogramming of host energy metabolism for pathogenesis. Front Cell Infect Microbiol 11:621156.

10. Jha HC, Banerjee S, Robertson ES. 2016. The role of gammaherpesviruses in cancer pathogenesis. Pathogens 5:18.

11. Wang Q, Shi B, Yang G, Zhu X, Shao H, Qian K, Ye J, Qin A. 2023. Metabolomic profiling of Marek’s disease virus infection in host cell based on untargeted LC-MS. Front Microbiol 14:1270762.

12. Manchester M, Anand A. 2017. Metabolomics: Strategies to define the role of metabolism in virus infection and pathogenesis. Adv Virus Res 98:57–81.

13. Diamond DL, Syder AJ, Jacobs JM, Sorensen CM, Walters K-A, Proll SC, McDermott JE, Gritsenko MA, Zhang Q, Zhao R, Metz TO, Camp DG 2nd, Waters KM, Smith RD, Rice CM, Katze MG. 2010. Temporal proteome and lipidome profiles reveal hepatitis C virus-associated reprogramming of hepatocellular metabolism and bioenergetics. PLoS Pathog 6:e1000719.

14. Hollenbaugh JA, Munger J, Kim B. 2011. Metabolite profiles of human immunodeficiency virus infected CD4+ T cells and macrophages using LC–MS/MS analysis. Virology 415:153–159.

15. Birungi G, Chen SM, Loy BP, Ng ML, Li SFY. 2010. Metabolomics approach for investigation of effects of dengue virus infection using the EA.hy926 cell line. J Proteome Res 9:6523–6534.

16. Vastag L, Koyuncu E, Grady SL, Shenk TE, Rabinowitz JD. 2011. Divergent effects of human cytomegalovirus and herpes simplex virus-1 on cellular metabolism. PLoS Pathog 7:e1002124.

17. Munger J, Bajad SU, Coller HA, Shenk T, Rabinowitz JD. 2006. Dynamics of the cellular metabolome during human cytomegalovirus infection. PLoS Pathog 2:e132.

18. Wang LW, Shen H, Nobre L, Ersing I, Paulo JA, Trudeau S, Wang Z, Smith NA, Ma Y, Reinstadler B, Nomburg J, Sommermann T, Cahir-McFarland E, Gygi SP, Mootha VK, Weekes MP, Gewurz BE. 2019. Epstein-Barr-virus-induced one-carbon metabolism drives B cell transformation. Cell Metab 30:539–555.e11.

19. Delgado T, Sanchez EL, Camarda R, Lagunoff M. 2012. Global metabolic profiling of infection by an oncogenic virus: KSHV induces and requires lipogenesis for survival of latent infection. PLoS Pathog 8:e1002866.

20. Delgado T, Carroll PA, Punjabi AS, Margineantu D, Hockenbery DM, Lagunoff M. 2010. Induction of the Warburg effect by Kaposi’s sarcoma herpesvirus is required for the maintenance of latently infected endothelial cells. Proc Natl Acad Sci U S A 107:10696–10701.

21. Sanchez EL, Pulliam TH, Dimaio TA, Thalhofer AB, Delgado T, Lagunoff M. 2017. Glycolysis, glutaminolysis, and fatty acid synthesis are required for distinct stages of Kaposi’s sarcoma-associated Herpesvirus lytic replication. J Virol 91.

22. Sychev ZE, Hu A, DiMaio TA, Gitter A, Camp ND, Noble WS, Wolf-Yadlin A, Lagunoff M. 2017. Integrated systems biology analysis of KSHV latent infection reveals viral induction and reliance on peroxisome mediated lipid metabolism. PLoS Pathog 13:e1006256.

23. Alfaez A, Christopher MW, Garrett TJ, Papp B. 2025. Analysis of metabolomic reprogramming induced by infection with Kaposi’s sarcoma-associated Herpesvirus using untargeted metabolomic profiling. Int J Mol Sci 26:3109.

24. Wan Q, Tavakoli L, Wang T-Y, Tucker AJ, Zhou R, Liu Q, Feng S, Choi D, He Z, Gack MU, Zhao J. 2024. Hijacking of nucleotide biosynthesis and deamidation-mediated glycolysis by an oncogenic herpesvirus. Nat Commun 15:1442.

25. Clark SA, Vazquez A, Furiya K, Splattstoesser MK, Bashmail AK, Schwartz H, Russell M, Bhark S-J, Moreno OK, McGovern M, Owsley ER, Nelson TA, Sanchez EL, Delgado T. 2023. Rewiring of the host cell metabolome and lipidome during lytic gammaherpesvirus infection is essential for infectious-virus production. J Virol 97:e0050623.

26. Brulois KF, Chang H, Lee AS-Y, Ensser A, Wong L-Y, Toth Z, Lee SH, Lee H-R, Myoung J, Ganem D, Oh T-K, Kim JF, Gao S-J, Jung JU. 2012. Construction and manipulation of a new Kaposi’s sarcoma-associated herpesvirus bacterial artificial chromosome clone. J Virol 86:9708–9720.

27. Cronstein BN, Aune TM. 2020. Methotrexate and its mechanisms of action in inflammatory arthritis. Nat Rev Rheumatol 16:145–154.

28. Bedoui Y, Guillot X, Sélambarom J, Guiraud P, Giry C, Jaffar-Bandjee MC, Ralandison S, Gasque P. 2019. Methotrexate an old drug with new tricks. Int J Mol Sci 20:5023.

29. Koźmiński P, Halik PK, Chesori R, Gniazdowska E. 2020. Overview of dual-acting drug methotrexate in different neurological diseases, autoimmune pathologies and cancers. Int J Mol Sci 21:3483.

30. Curreli F, Cerimele F, Muralidhar S, Rosenthal LJ, Cesarman E, Friedman-Kien AE, Flore O. 2002. Transcriptional downregulation of ORF50/Rta by methotrexate inhibits the switch of Kaposi’s sarcoma-associated herpesvirus/human herpesvirus 8 from latency to lytic replication. J Virol 76:5208–5219.

31. Chen J, Zhang H, Chen X. 2020. Pemetrexed inhibits Kaposi’s sarcoma-associated herpesvirus replication through blocking dTMP synthesis. Antiviral Res 180:104825.

32. Yang R, Li B. 2024. UDPG: Maintaining the true nature of sugar. Cell Mol Immunol 21:1087–1088.

33. Bird AR, Molloy JC, Hall EAH. 2023. Biocatalytic synthesis of 2’-deoxynucleotide 5’-triphosphates from bacterial genomic DNA: Proof of principle. Biotechnol Bioeng 120:1531–1544.

34. Dunn J, Grider MH. 2025. Physiology, adenosine triphosphateStatPearls. StatPearls Publishing, Treasure Island (FL).

35. Menezo Y, Elder K, Clement A, Clement P. 2022. Folic acid, folinic acid, 5 methyl TetraHydroFolate supplementation for mutations that affect epigenesis through the folate and one-carbon cycles. Biomolecules 12:197.

36. Fontaine KA, Camarda R, Lagunoff M. 2014. Vaccinia virus requires glutamine but not glucose for efficient replication. J Virol 88:4366–4374.

37. Qin Z, Freitas E, Sullivan R, Mohan S, Bacelieri R, Branch D, Romano M, Kearney P, Oates J, Plaisance K, Renne R, Kaleeba J, Parsons C. 2010. Upregulation of xCT by KSHV-encoded microRNAs facilitates KSHV dissemination and persistence in an environment of oxidative stress. PLoS Pathog 6:e1000742.

38. Strahan RC, McDowell-Sargent M, Uppal T, Purushothaman P, Verma SC. 2017. KSHV encoded ORF59 modulates histone arginine methylation of the viral genome to promote viral reactivation. PLoS Pathog 13:e1006482.

39. Tankersley RW Jr. 1964. Amino acid requirements of herpes simplex virus in human cells. J Bacteriol 87:609–613.

40. Zhang T, Wang Y, Zhang L, Liu B, Xie J, Wood C, Wang J. 2011. Lysine residues of interferon regulatory factor 7 affect the replication and transcription activator-mediated lytic replication of Kaposi’s sarcoma-associated herpesvirus/human herpesvirus 8. J Gen Virol 92:181–187.

41. Rhodes DA, Boyle LH, Boname JM, Lehner PJ, Trowsdale J. 2010. Ubiquitination of lysine-331 by Kaposi’s sarcoma-associated herpesvirus protein K5 targets HFE for lysosomal degradation. Proc Natl Acad Sci U S A 107:16240–16245.

42. González CM, Wang L, Damania B. 2009. Kaposi’s sarcoma-associated herpesvirus encodes a viral deubiquitinase. J Virol 83:10224–10233.

43. Sanchez EL, Carroll PA, Thalhofer AB, Lagunoff M. 2015. Latent KSHV Infected Endothelial Cells Are Glutamine Addicted and Require Glutaminolysis for Survival. PLoS Pathog 11:e1005052.

44. Zhu Y, Li T, Ramos da Silva S, Lee J-J, Lu C, Eoh H, Jung JU, Gao S-J. 2017. A critical role of glutamine and asparagine γ-nitrogen in nucleotide biosynthesis in cancer cells hijacked by an oncogenic virus. MBio 8.

45. Fan Q, Longnecker R, Connolly SA. 2021. Herpes simplex virus glycoprotein B mutations define structural sites in domain I, the membrane proximal region, and the cytodomain that regulate entry. J Virol 95:e0105021.

46. Karthigeyan KP, Connors M, Binuya CR, Gross M, Fuller AS, Crooks CM, Wang H-Y, Sponholtz MR, Byrne PO, Herbek S, Andy C, Gerber LM, Campbell JD, Williams CA, Mitchell E, van der Maas L, Miller I, Yu D, Bottomley MJ, McLellan JS, Permar SR. 2025. A human cytomegalovirus prefusion-like glycoprotein B subunit vaccine elicits humoral immunity similar to that of postfusion gB in mice. J Virol 99:e0217824.

47. Ito F, Zhen J, Xie G, Huang H, Silva JC, Wu T-T, Zhou ZH. 2025. Structure of the Kaposi’s sarcoma-associated herpesvirus gB in post-fusion conformation. J Virol 99:e0153324.

48. He M, Zhou X, Wang X. 2024. Glycosylation: mechanisms, biological functions and clinical implications. Signal Transduct Target Ther 9:194.

49. Mortazavi Y, Lidenge SJ, Tran T, West JT, Wood C, Tso FY. 2020. The Kaposi’s sarcoma-associated Herpesvirus (KSHV) gH/gL complex is the predominant neutralizing antigenic determinant in KSHV-infected individuals. Viruses 12:256.

50. Zhu FX, Chong JM, Wu L, Yuan Y. 2005. Virion proteins of Kaposi’s sarcoma-associated herpesvirus. J Virol 79:800–811.

51. Neipel F, Albrecht JC, Fleckenstein B. 1997. Cell-homologous genes in the Kaposi’s sarcoma-associated rhadinovirus human herpesvirus 8: determinants of its pathogenicity? J Virol 71:4187–4192.

52. Subramanian R, Sehgal I, D’Auvergne O, Kousoulas KG. 2010. Kaposi’s sarcoma-associated herpesvirus glycoproteins B and K8.1 regulate virion egress and synthesis of vascular endothelial growth factor and viral interleukin-6 in BCBL-1 cells. J Virol 84:1704–1714.

53. DeVito SR, Ortiz-Riaño E, Martínez-Sobrido L, Munger J. 2014. Cytomegalovirus-mediated activation of pyrimidine biosynthesis drives UDP-sugar synthesis to support viral protein glycosylation. Proc Natl Acad Sci U S A 111:18019–18024.

54. Nandakumar D, Glaunsinger B. 2019. An integrative approach identifies direct targets of the late viral transcription complex and an expanded promoter recognition motif in Kaposi’s sarcoma-associated herpesvirus. PLoS Pathog 15:e1007774.

55. Purushothaman P, Uppal T, Verma SC. 2015. Molecular biology of KSHV lytic reactivation. Viruses 7:116–153.

56. DHFR dihydrofolate reductase [Homo sapiens (human)] - Gene - NCBI. https://www.ncbi.nlm.nih.gov/gene/1719. Retrieved 17 November 2025.

57. Cinquina CC, Grogan E, Sun R, Lin SF, Beardsley GP, Miller G. 2000. Dihydrofolate reductase from Kaposi’s sarcoma-associated herpesvirus. Virology 268:201–217.

58. Furst DE, Kremer JM. 1988. Methotrexate in rheumatoid arthritis. Arthritis Rheum 31:305–314.

59. Stenger AA, Houtman PM, Bruyn GA. 1992. Does folate supplementation make sense in patients with rheumatoid arthritis treated with methotrexate? Ann Rheum Dis 51:1019–1020.

60. Russo JJ, Bohenzky RA, Chien MC, Chen J, Yan M, Maddalena D, Parry JP, Peruzzi D, Edelman IS, Chang Y, Moore PS. 1996. Nucleotide sequence of the Kaposi sarcoma-associated herpesvirus (HHV8). Proc Natl Acad Sci U S A 93:14862–14867.

61. Liang JH, Wang C, Yiu SPT, Zhao B, Guo R, Gewurz BE. 2021. Epstein-Barr virus induced cytidine metabolism roles in transformed B-cell growth and survival. MBio 12:e0153021.

62. Myoung J, Ganem D. 2011. Generation of a doxycycline-inducible KSHV producer cell line of endothelial origin: maintenance of tight latency with efficient reactivation upon induction. J Virol Methods 174:12–21.

