## Supplementary figures and images for "Global Metabolomic Analysis of Lytic KSHV Infection: Induced Host Nucleotide Metabolism is Required for Infectious Virus Production"

### Supplemental Figure 1

#### Supplementary Figure 1

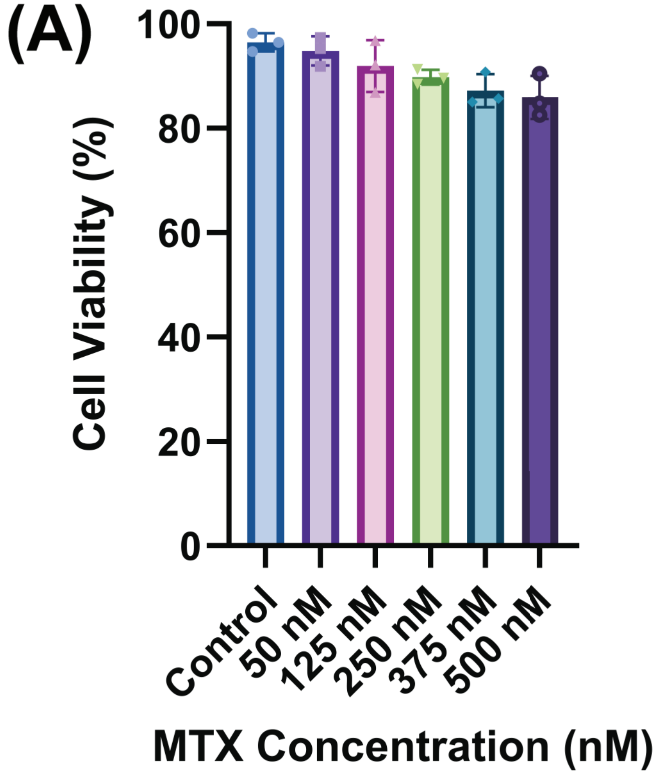
